# PhyloSophos: a high-throughput scientific name mapping algorithm augmented with explicit consideration of taxonomic science

**DOI:** 10.1101/2023.03.17.533059

**Authors:** Min Hyung Cho, Kwang-Hwi Cho, Kyoung Tai No

**Affiliations:** Bioinformatics and Molecular Design Research Center (BMDRC). 209, Veritas A Hall, Yonsei University, 85 Songdogwahak-ro, Yeonsu-gu, Incheon 21983, Republic of Korea; School of Systems Biomedical Science and Research Center for Integrative Basic Science, Soongsil University, Seoul 06978, South Korea; Department of Integrative Biotechnology & Translational Medicine, Yonsei University. 214, Veritas A Hall, Yonsei University, 85 Songdogwahak-ro, Yeonsu-gu, Incheon 21983, Republic of Korea

## Abstract

**Summary:** The nature of taxonomic science and the scientific nomenclature system makes it difficult to use scientific names as identifiers without running into complications. To facilitate high-throughput analysis of biological data involving scientific names, we designed PhyloSophos, a Python package that takes into account the properties of scientific names and taxonomic systems to map name inputs to the entries within the reference database of choice. We would like to present three case-studies which demonstrates how our implementations, including rule-based pre-processing and recursive mapping could improve mapping performance and information availability. We expect PhyloSophos to help with the systematic processing of poorly digitized and curated biological data, such as biodiversity information and ethnopharmacological resources, thus enabling full-scale bioinformatics analysis using these data.

**Availability and implementation:** PhyloSophos is available at GitHub https://github.com/mhcho4096/phylosophos.

**Supplementary information:** Supplementary data are available at Bioinformatics online.

## [1] Introduction

While scientific names are accepted and used as universal identifiers for biological species, they are inherently poor identifiers for several reasons (Remsen et al., 2016). Scientific names lack certain desirable qualities of an ideal database identifier (McMurry et al. 2017), such as stability, uniqueness, persistence, and convertibility (Mozzherin et al. 2017; Thiele et al. 2021; Thomson et al. 2021), thus making the process of verifying and standardizing scientific names across multiple resources complex.

To address this challenge, multiple algorithmic approaches such as Taxamatch (Rees, 2014) and gnparser (Mozzherin et al. 2017) have been developed to facilitate the process of recognizing, correcting, and managing scientific names. Taxamatch incorporates a modified Damerau-Levenshtein distance and phonetic algorithm to detect errors within scientific names, thus functioning as a high-throughput scientific name processing pipeline. On the other hand, gnparser identifies semantic elements within scientific names such as abbreviations and authorship, and then processes them to provide the canonical form of the given input. These applications can be utilized to handle tasks related to scientific names, such as curating biodiversity information (Montgomery et al. 2019; Conti et al. 2021), thereby facilitating research that would otherwise require significant time to manually curate the data.

Here, we would like to introduce PhyloSophos, a high-throughput scientific name processor designed to provide connections between scientific name inputs and a specific taxonomic system. PhyloSophos is conceptually a mapper that returns the corresponding taxon identifier from a reference of choice: to maximize performance, PhyloSophos can refer to multiple available references to search for synonyms and recursively map them into a chosen reference. It also corrects common Latin variants and vernacular names, which often appear in ethnobotanical literature and natural products research (Bennett and Balick, 2014; Dauncey et al. 2016), subsequently returns proper scientific names and its corresponding taxon identifiers. We would like to provide three case-studies which demonstrate mapping scientific names found in three NP databases, thereby represent the capacity of PhyloSophos to process scientific name variants with superior performance.

## [2] Implementation

PhyloSophos is a Python-based standalone package that could be divided into four parts: taxonomic reference setup, input pre-processing, scientific name mapping, and data export (Figure 1A).

**Figure 1.**
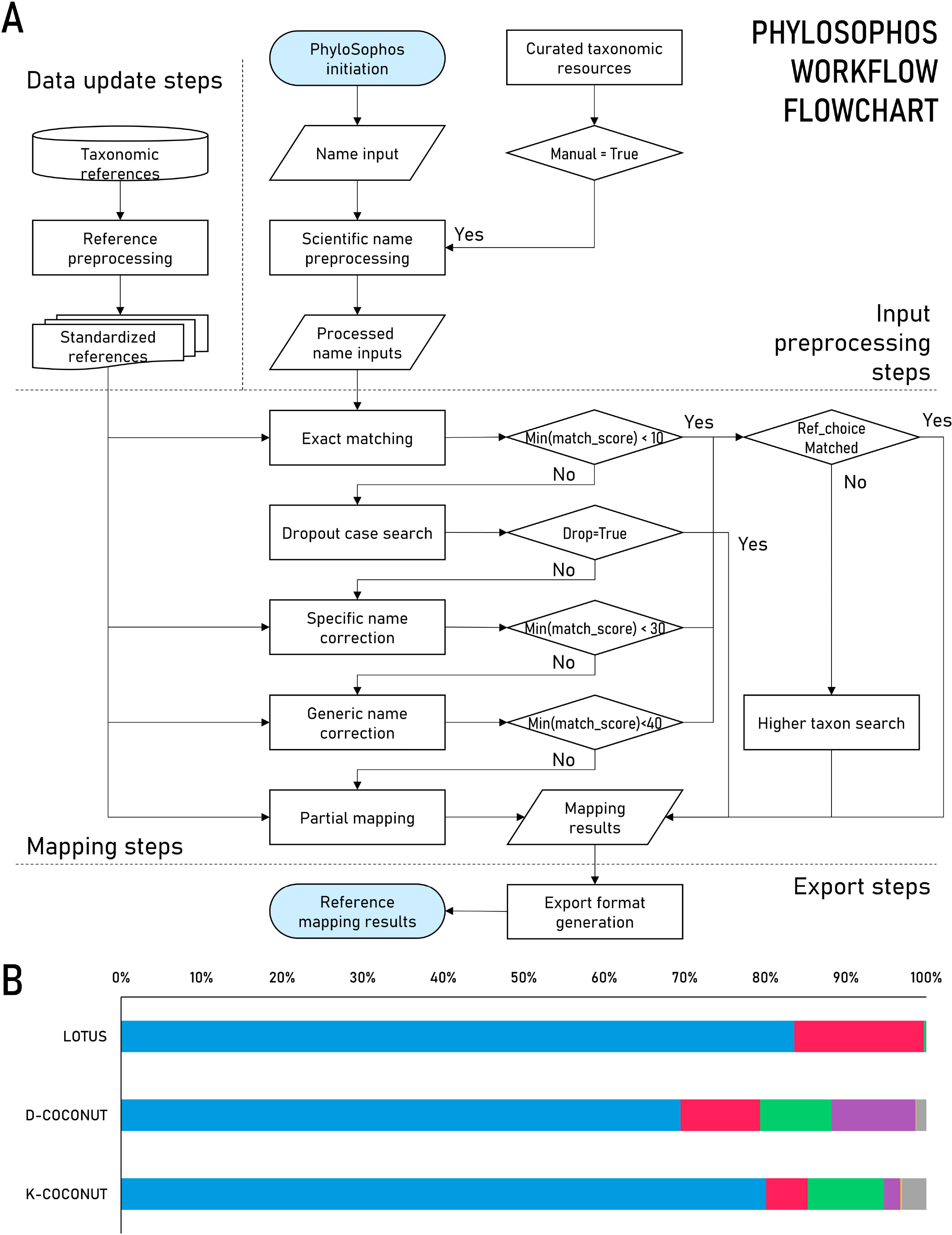
A: Flowchart of PhyloSophos. B: Mapping statuses of scientific names found within three case-study databases (see supplementary table 2/3). Blue: Valid name & exact match, Red: Valid name & nearest match, Green: Corrected match, Purple: Partial match, Yellow: Dropout (mapping denied) cases, Gray: Unmatched.

Taxonomic reference setup is done by executing the separate update script, which enables the user to automatically download and incorporate up-to-date taxonomic information from three databases, Catalogue of Life (Bánki et al. 2022), Encyclopedia of Life (Parr et al. 2014), and NCBI taxonomy (Schoch et al. 2020). It generates two files, one of which lists the first word-blocks of the scientific names found within the reference, along with associated DB identifiers, and the other lists all taxonomic entities along with their synonyms, taxonomic ranks, and phylogenetic lineage. The user may further customize their own PhyloSophos environment by placing additional taxonomic reference data files within the reference directory (Supplementary text S4). We provide an example script of including extra taxonomic resources using the GBIF Backbone Taxonomy, which could be utilized as a template for generating update scripts for other references.

Input pre-processing involves several distinct sub-modules to optimize mapping performance (Supplementary text S2). If the query does not match any of the taxonomic names found in the incorporated references, a semantic rule-based screening is performed to identify queries that should be treated differently or excluded before the mapping step. For example, if abbreviations like ‘cf.’ or ‘aff.’ is found within a query, which denotes a similarity to a given taxon (but not that taxon exactly), PhyloSophos first removes the abbreviations first, tries to match it, and then returns an identifier of a higher taxon which is found within the phylogenetic lineage of the match, to reflect the connotation of given abbreviation. Similarly, if the query includes certain word-blocks such as ‘phytoplasma’ or ‘endosymbiont’, the query is ignored, as the scientific name included within the query, which is mostly that of the host species, does not convey taxonomic information about the query itself.

Correction of Latin inflection is deployed to properly recognize scientific names in variant forms, which is often found in traditional medicine-related resources. This step removes word-blocks that are likely not part of scientific names, such as those denoting specific parts of a species, and reconstructs the most likely original form of the given query, which can then be mapped to find a match within a reference. PhyloSophos can also take a manually curated list of [vernacular name-scientific name] pairs and apply it during the pre-processing step (Supplementary text S5), which allows for further translation of vernacular names before (most likely futile) the mapping step and thus expedites the processing.

Scientific name mapping is done reference-wise with Damerau-Levenshtein distance algorithm (Wagner and Lowrance, 1975), and then recursive mapping - search of scientific names and their synonyms from other references within the chosen taxonomic reference - is done to search whether a better mapping is possible with information from other references. Identified mapping results are returned as a table along with its mapping status (Supplementary text S6), along with all other ‘best matches’ found in other databases: this provides information as to whether the individual query is a legitimate scientific name which is just not included in the chosen reference, or if it is not likely a scientific name at all.

## [3] Case studies

### [3-1] Mapping of scientific names found in LOTUS initiative

LOTUS initiative is an open database which includes more than 750,000+ species-compound pairs along with their corresponding references (Rutz et al. 2022). After removing duplicate entries, we identified 669,025 species-compound pairs from the LOTUS metadata, which involves 38,173 different species. The metadata also provided taxonomic reference database IDs for each species, including NCBI taxonomy ID for 28,839 species and GBIF taxon identifier for 33,869 species (Supplementary data SD1, see also Supplementary text S6).

We used PhyloSophos to analyze LOTUS species entries and investigate whether our algorithm could improve this database. Thanks to rigorous harmonization efforts, 38,044 names found in LOTUS metadata was identified as a valid scientific name. Interestingly, we found multiple species entries, which were not mapped to either NCBI taxonomy of GBIF, could be precisely linked to taxonomic entities within the references (Figure 1B, SD1). PhyloSophos provided 31,915 species with an exact match to one of NCBI’s taxonomy entries, and 36,282 species with that to one of GBIF’s entries, leaving only 1 species entirely unmapped. We identified 36 cases of disagreement between the PhyloSophos mapping results and the NCBI IDs provided in the original metadata, and individual manual curation identified most of the disagreements that arose due to periodic updates of taxonomic references (SD1).

This ultimately reduces the number of species-compound pairs without either NCBI or GBIF taxonomic reference from 34,695 to 5,560 (SD1), suggesting that PhyloSophos could maximize the utility of the LOTUS database by salvaging as much information from the metadata as possible.

### [3-2] Mapping of scientific names found in COCONUT

The Collection of Open Natural Products (COCONUT) database is one of the largest natural products (NP) database which was assembled from at least 53 different data sources (Sorokina et al. 2021). While COCONUT provides 407,270 unique NP entries, only 83,264 of them were annotated with associated species: we were able to identify 28,528 species and 377,373 species-compound pairs from these entries (Supplementary data SD2).

We conducted a similar analysis with PhyloSophos to investigate the scientific names found in the COCONUT database (we will refer it D-COCONUT), and found a significantly larger number of names that were problematic (SD2). Only 19,823 names could be exactly mapped with NCBI taxonomy, and further 319 names could be mapped at the species-level (Supplementary tables 1 & 2). Among these mapped entries, 2,265 of these entries were mapped recursively using sub-results generated using other taxonomic databases. We could also identify names containing typographical errors, which often appear in scientific names (Patterson et al. 2016): 2,525 name inputs were mapped to the correct taxonomic entity using an edit distance-based algorithm (Figure 1B).

An additional 5,799 names were mapped at the higher taxonomic level, and 368 names were left unmapped (Supplementary table 2). We identified multiple scientific names in the input list that were without proper specific epithets but were accompanied by strain names: as PhyloSophos only accepts exact matches for strain-level mapping, edit distance-based search was not applied in these cases, resulting in many name inputs being mapped to higher-level taxonomic entities.

Further screening and analysis revealed that 318,376 species-compound pairs were mapped with NCBI taxonomy, at least at the genus-level (SD2). Meaningful species-compound pairs were significantly lower, as the significant fraction of pairs were associated with non-descriptive terms such as ‘plant’, ‘eukaryote’, or ‘marine’, rendering it practically unusable in precise analysis.

### [3-3] Mapping of scientific names with Latin inflections

The Compound Combination-Oriented Natural Product Database (which interestingly shares the abbreviation, COCONUT: we will refer it K-COCONUT) is a database that not only contains entities related to natural products, but also other biological information such as genes, proteins, and disorders (Yoo et al. 2018). We acquired raw data from the previously available database website, which encompasses 8,642 ‘herbs’: these include not only plants but also fungi, animals, and even non-biological entities (Supplementary data SD3). We identified at least 334 names affected by Latin inflection (SD3) and found that PhyloSophos could successfully extract putative ‘original’ forms from the names, perform edit distance-based searches, and ultimately provide taxonomic identifiers for each case (Supplementary tables 1&2, SD3).

We are aware that traditional names often convey incomplete information about the biological identity of itself, for example by omitting the specific epithet: still, PhyloSophos could be utilized to extract the maximum available taxonomic information from the given query, such as genus-level information.

## [4] Discussion

We have demonstrated that PhyloSophos can systematically map scientific name inputs into the corresponding taxonomic entities within the reference database of choice. By considering the properties of taxonomy and nomenclature systems, we were able to apply several ideas such as rule-based preprocessing and multiple reference-based recursive mapping to the application, resulting in increased resolving power. Despite the fact that there is still room for optimization, such as reducing memory usage, PhyloSophos is already capable of delivering reliable results for practical research. We expect this application will help facilitate the systematic incorporation of disparate biological data, such as biodiversity reports and ethnobotanical annotations, into the knowledge network, thereby providing access to high-throughput analysis.

## Acknowledgements

This research was funded and supported by Korea Institute for Advancement of Technology (KIAT) of the Republic of Korea (Project name: Establishment and Demonstration of bio-material data platform) (Project number: P0014714).

