## Supplementary material for "PhyloSophos: a high-throughput scientific name mapping algorithm augmented with explicit consideration of taxonomic science": supplementary_material_phylosophos.docx

**Contents**

**Supplementary texts	…	2**

Text S1. Short description	…	2

Text S2. Detailed PhyloSophos workflow	…	2

Text S3. Mapping status codes & description	…	4

Text S4. PhyloSophos modding guide: incorporation of new reference	…	8

Text S5. PhyloSophos modding guide: incorporation of manual curation data	…	9

Text S6. Case study details & supplementary data files description	…	10

**Supplementary figures**	…	11

Supplementary figure 1. Main function of PhyloSophos core mapper script	…	11

Supplementary figure 2. Pseudocode of mapper initialization function	…	12

Supplementary figure 3. Pseudocode of scientific name input import function	…	13

Supplementary figure 4. Pseudocode of input file-wise PhyloSophos mapping function	…	14

Supplementary figure 5. Pseudocode of PhyloSophos intrageneric correction subfunction	…	15

**Supplementary tables**	…	16

Supplementary table 1. PhyloSophos mapping status for scientific names found in natural
products databases used in the case study	…	16

Supplementary table 2. PhyloSophos broad mapping status (count)	…	17

Supplementary table 3. PhyloSophos broad mapping status (ratio)	…	17

**Supplementary references**	…	18

**Supplementary texts**

**Text S1. Short description**

PhyloSophos is a high-throughput scientific name processor which achieves greater mapping performance by referencing multiple taxonomic references and recognizing the semantic structure of scientific names. It also corrects common Latin variants and vernacular names, which often appear in various biological databases and resources.

PhyloSophos is a Python-based standalone package: it takes one or more files with a list of putative scientific names as an input and returns a machine-readable table which shows comprehensive mapping information, thereby facilitating the use of scientific names in various types of data analysis. We anticipate this application to aid in the integration of a range of biological data, like biodiversity reports and ethnobotanical annotations, into the knowledge network we are currently accessible.

License information and detailed user guide for PhyloSophos could be found in the Github repository (https://github.com/mhcho4096/phylosophos).

**Text S2. Detailed PhyloSophos workflow**

PhyloSophos is a Python-based standalone package that could be divided into four parts: taxonomic reference setup, input pre-processing, scientific name mapping, and data export (See main figure 1A). The main function, which appear in 'phylosophos_core_py' script, is shown (Supplementary figure 1). The PhyloSophos core mapper begins by recognizing optional arguments from the command and modifying the parameters as necessary (Supplementary figure 2). It recognizes five types of optional argument types.

- Help (-h, -help, -guide): if one of these arguments is given, a hard-coded guide to PhyloSophos will appear in the console. This will provide simple instructions on how to customize PhyloSophos mapping parameters. No following parameter value is required.

- Reference type change (-r, -ref): if one of these arguments is given, PhyloSophos will change the database of choice to the one specified by the following argument. The default setting is 'ncbi', while 'col' and 'eol' are also available in basic PhyloSophos system.

- Input type change (-i, -input): if one of these arguments is given, along with the name of the input file, PhyloSophos will specifically import the given file as an input. If not (as a default setting), PhyloSophos will consider all files within the '/input' directory to be scientific name input files.

- Levenshtein distance cutoff (-l, -lev, -cutoff): if one of these arguments is given, along with an integer value, PhyloSophos will change the edit distance cutoff (default setting = 3) to the specified value.

- Manual curation status (-m, -manual, -curation): if one of these arguments is given, along with a value 1, PhyloSophos will import '/pp_learning/manual_curation_list.tsv' and utilize this information to pre-process inputs. If not (as a default setting), PhyloSophos will not import extra information other than reference data files within '/pp_ref' directory.

After the initialization process, PhyloSophos begins importing scientific name input files (Supplementary figure 3). Depending on the input type parameter, it may import every file within the '/input' directory (default setting) or a specified single input file. After this step, PhyloSophos imports formatted reference files from the '/pp ref' directory, generating four taxonomic dictionaries.

- tax_ref_list: dictionary of available taxonomic reference types (e.g. 'col', 'eol', 'ncbi')

- ref_names_dict: dictionary of all taxonomic entities found within each taxonomic reference

- ref_genus_dict: dictionary of all first word-blocks (mostly generic epithets, synonyms included) found within each taxonomic reference, and list of taxonomic entities associated with given word-block

- ref_raw_names: dictionary of all unique scientific names and synonyms (partly overlaps with ref_genus_dict) found within each taxonomic reference, and list of entities associated with given name

Based on the taxonomic references, the PhyloSophos mapping function examines each scientific name input and maps it to the corresponding taxonomic entities (Supplementary figure 4). The workflow can be described as follows.

- Exact matching: PhyloSophos first attempts to find an exact match with a given string, either as-is or after making a simple correction. If a match is found in a particular reference, it is given a status code 0~5 (depending on the exact mapping status) for that reference. If a match is found in one or more references but not in others, PhyloSophos collects synonym information from the matched taxonomic entity and searches for those in the references without an exact match (recursive search). If a match is found with a recursive search, it is given a status code 6 or 8.

- Nearest taxon mapping: If the given input is exactly mapped to a taxonomic entity in at least one taxonomic reference, PhyloSophos considers the input is a valid scientific name and does not consider it a target for edit distance-based correction. Instead, it extracts phylogenetic lineage information from the previously matched taxonomic entity and attempts to provide the nearest taxon (with the lowest taxonomic rank) for references without an exact match. It is given a status code 10~18, depending on the taxonomic rank of provided taxon.

- Rule-based input dropout: Before calling edit distance-based correction methods, PhyloSophos first checks whether the given input has a particular keyword and should not be processed further (see text S3). It is given a status code 90~99, depending on the keyword the input contains.

- Edit-distance based correction (specific epithet only): As Damerau-Levenshtein edit distance calculation requires significant computing resources, PhyloSophos attempts to search within a limited pool of scientific names first (Supplementary figure 5). It first searches for scientific names with the same first word-block, then further reduces the pool using the length and composition of characters found within each string. Calculation of edit distance follows, and the scientific name with the lowest edit distance is returned as the taxonomic entity corresponding to the given input (along with a status code 20~24, depending on the mapping status).

- Edit-distance based correction (whole input): If a mapping is not possible with previous steps, similar edit distance-based correction using whole string, along with Latin inflection correction, is attempted. It is given a status code 30~36, depending on the mapping status.

- Partial mapping: If a mapping is not possible with previous steps, PhyloSophos attempts to provide mapping results using a part of given inputs. If a partial mapping is possible, it is given a status code 100, and if not, it is given a status code 1000.

Calculated mapping results are compiled to a result file (per each input file) and exported to '/result' directory. The name of the result file will be 'phylosophos_result_[export_date]_[export_time]_[input_file_name]'. [export_date] and [export_time] will be six-digit numbers.

**Text S3. Mapping status codes & description**

Detailed descriptions of each mapping status codes are as follows.

● Codes 0~9: Valid scientific name/Exact match.

- Code 0: Exact match found within a DB of choice / No correction / matched with a canonical name

- Input is found within a DB of choice as-is. It is matched with canonical name of a taxonomic entity.

- Code 1: Exact match found within a DB of choice / No correction / matched with a single synonym

- Input is found within a DB of choice as-is. It is matched with synonym of a taxonomic entity.

- Code 2: Exact match found within a DB of choice / No correction / matched with multiple entities

- Input is found within a DB of choice as-is. It is matched with multiple taxonomic entities.

- It is possible that a single scientific name, without proper authority information, could refer to multiple entities within a taxonomic database. Most of these cases involve single-word input and fall into one of three categories:

- Genus and subgenus: Many subgenera share the same epithet as the genera they are included in (e.g. subgenus Rhododendron within genus Rhododendron).

- Taxa in different phylogenetic domains: Scientific names are governed by multiple nomenclature codes, which govern specific types of taxonomic groups (e.g. animals, plants, bacteria). As these nomenclature codes are independent of one another, it is possible that a single generic epithet is used for multiple genera in different taxonomic groups (e.g. *Anisoptera*, *Callicarpa*, *Darwinella*).

- Obsolete synonyms: There were several instances where a single scientific name was applied to multiple taxa (e.g. *Polyporus badius*). As it is a violation of the nomenclature code, misapplied names are later corrected by taxonomic authorities; however, those names remain in a record of obsolete synonyms. If a given input name is not found in a canonical list of scientific names but is found multiple times in a synonym list, PhyloSophos will return all entities associated with that name.

- Manual review is highly recommended in these cases.

- Code 3: Exact match found within a DB of choice / Simple correction / matched with a canonical name

- Input is found within a DB of choice after a simple correction process. (similar to code 0) It is matched with canonical name of a taxonomic entity.

- Code 4: Exact match found within a DB of choice / Simple correction / matched with a single synonym

- Input is found within a DB of choice after a simple correction process. (similar to code 1) It is matched with synonym of a taxonomic entity.

- Code 5: Exact match found within a DB of choice / Simple correction / matched with multiple entities

- Input is found within a DB of choice after a simple correction process. (similar to code 2) It is matched with multiple taxonomic entities.

- Code 6: Exact match found within other DBs / Synonym information is used to search in a DB of choice / Matched with a single entity

- Input is found in more than one of the other taxonomic databases, but not in the DB of choice. To identify the corresponding taxonomic entity within a DB of choice, we collected synonym information from other databases and searched for these names again in a DB of choice. As a result, (similar to codes 0,1) a single taxonomic entity was identified.

- Code 8: Exact match found within other DBs / Synonym information is used to search in a DB of choice / Matched with multiple entities

- Input is found in more than one of the other taxonomic databases, but not in the DB of choice. To identify the corresponding taxonomic entity within a DB of choice, we collected synonym information from other databases and searched for these names again in a DB of choice. As a result, (similar to code 2) multiple taxonomic entities were identified.

● Codes 10~19: Valid scientific name / Nearest taxon match.

- If a given input is mapped to a taxonomic entity with a mapping code of less than 10, PhyloSophos considers it to be a valid scientific name and does not consider it a target for edit distance-based correction. If a given input is not found within a DB of choice, nearest taxon match algorithm is applied instead.

- PhyloSophos extracts phylogenetic lineage information from a taxonomic entity that corresponds to a given input. Starting from the lowest taxonomic rank, the algorithm searches for the taxonomic entity within a DB of choice that exactly matches (as code 0) the name of the higher-rank taxon. This method allows to identify the lowest (‘nearest’) taxonomic entity which includes the given input as a member.

- Different codes are applied based on the taxonomic level of the matched taxon.

- Code 10: Nearest match to the species level

- Code 11: Nearest match to the genus level

- Code 12: Nearest match to the family level

- Code 13: Nearest match to the order level

- Code 14: Nearest match to the class level

- Code 15: Nearest match to the phylum level

- Code 16: Nearest match to the kingdom level

- Code 17: Nearest match to the domain level

● Codes 20~29: Specific epithet corrected.

- Code 20: Specific epithet corrected / Exact match found within a DB of choice / Matched with a single entity

- Corrected input is found within a DB of choice. It is matched with a single taxonomic entity within a DB.

- Code 21: Specific epithet corrected / Exact match found within a DB of choice / Matched with multiple entities

- Corrected input is found within a DB of choice. It is matched with multiple taxonomic entities within a DB.

- Code 22: Specific epithet corrected / Exact match found within other DBs / Synonym information is used to search in a DB of choice / Matched with a single entity

- Corrected input is found in more than one of the other taxonomic databases, but not in the DB of choice. To identify the corresponding taxonomic entity within a DB of choice, we collected synonym information from other databases and searched for these names again in a DB of choice. As a result, a single taxonomic entity was identified.

- Code 23: Specific epithet corrected / Exact match found within other DBs / Synonym information is used to search in a DB of choice / Matched with multiple entities

- Corrected input is found in more than one of the other taxonomic databases, but not in the DB of choice. To identify the corresponding taxonomic entity within a DB of choice, we collected synonym information from other databases and searched for these names again in a DB of choice. As a result, multiple taxonomic entities were identified.

- Code 24: Specific epithet corrected / Exact match found within other DBs / Nearest taxon match

- If a corrected input is mapped to a taxonomic entity with a mapping code of less than 24 in one of the other databases, but not in the DB of choice, the nearest taxon match algorithm is applied (see codes 10~19).

● Codes 30~39: Generic/specific epithet corrected.

- Code 30: Latin inflection corrected / Exact match found within a DB of choice

- One of the possible 'original forms' of a given input is found within a DB of choice.

- Code 31: Latin inflection corrected / Exact match found within other DBs

- One of the possible 'original forms' of a given input is found in more than one of the other taxonomic databases, but not in the DB of choice. This code applies to both synonym-match (as code 3) and nearest taxon match (as code 24).

- Code 32: Generic/Specific epithet corrected / Exact match found within a DB of choice / Matched with a single entity

- Corrected input is found within a DB of choice. It is matched with a single taxonomic entity within a DB.

- Code 33: Generic/Specific epithet corrected / Exact match found within a DB of choice / Matched with multiple entities

- Corrected input is found within a DB of choice. It is matched with multiple taxonomic entities within a DB.

- Code 34: Generic/Specific epithet corrected / Exact match found within other DBs / Synonym information is used to search in a DB of choice / Matched with a single entity

- Corrected input is found in more than one of the other taxonomic databases, but not in the DB of choice. To identify the corresponding taxonomic entity within a DB of choice, we collected synonym information from other databases and searched for these names again in a DB of choice. As a result, a single taxonomic entity was identified.

- Code 35: Generic/Specific epithet corrected / Exact match found within other DBs / Synonym information is used to search in a DB of choice / Matched with multiple entities

- Corrected input is found in more than one of the other taxonomic databases, but not in the DB of choice. To identify the corresponding taxonomic entity within a DB of choice, we collected synonym information from other databases and searched for these names again in a DB of choice. As a result, multiple taxonomic entities were identified.

- Code 36: Generic/Specific epithet corrected / Exact match found within other DBs / Nearest taxon match

- If a corrected input is mapped to a taxonomic entity with a mapping code of less than 36 in one of the other databases, but not in the DB of choice, the nearest taxon match algorithm is applied (see codes 10~19).

● Codes 40~49: Correction denied.

- Code 40: Strain name involved / Nearest match.

- Strain codes are often very similar to one another, but they do not reflect the phylogenetic relationship between individual strains. Therefore, edit distance-based correction approaches could suggest scientific names with strain information, which are almost identical string-wise but not accurate in a taxonomic sense. If word-blocks containing strain-specific information (usually numeric characters) are identified within an input string, PhyloSophos removes them from the correction process and maps them to the nearest higher taxon (e.g. species).

- Code 41: Similarity-related abbreviation identified / Nearest match.

- Abbreviations such as "cf.", "aff.", and "sp.nov." denote similarity to the previously described taxon, but not necessarily identity. Therefore, it may be inappropriate to drop this abbreviation and revert to the scientific name of a taxon that is similar to the species which the input name describes. Instead, PhyloSophos first removes the abbreviations first, tries to match it, and then returns an identifier of a higher taxon which is found within the phylogenetic lineage of the match, to reflect the connotation of given abbreviation.

● Codes 90~99: Mapping denied.

- If a given input is not found within one of the databases as-is, and if it contains word-blocks that require special attention, then the further mapping process is denied and one of these codes is returned.

- Code 90: Non-organism flags

- Code 91: Unclassified-Uncultured-Unidentified

- Code 92: Environmental samples

- Code 93: Virus or phage

- Code 94: Phytoplasma

- Code 95: (endo)symbiont

- Code 96: Unresolvable hybrid

- Code 97: Multiple materia medica

- Manual review is highly recommended in these cases.

● Code 100: Partial mapping.

- If taxonomic mapping is not possible with a full input string, PhyloSophos attempts to find a taxonomic entity which matches a part of the input. It consecutively removes a word-block from the right side of the input string and searches for an exact match: if an exact match is found, the search process is terminated and a result is returned.

- It usually ends with genus or species level mapping (but not always).

● Code 1000: Unmapped input.

- For inputs that cannot be assigned to a corresponding taxonomic entity (not even partially), they are given the code.

**Text S4. PhyloSophos modding guide: incorporation of new reference**

PhyloSophos extracts two files from each taxonomic database and uses them as references. Each file has the following format (delimited by tabs).

● [ref]_node_dict.txt: [reference_identifier] - [reference_entity_canonical_name] - [reference_entity_synonyms] - [reference_entity_taxonomic_rank] - [reference_entity_phylogenetic_lineage] - [reference_entity_lineage_taxonomic_rank]

- reference_identifier: accession numbers of given entity within a database.

- reference_entity_canonical_name: canonical scientific name which the taxonomic database suggests for the given taxonomic entity.

- reference_entity_synonyms: alternative names which the taxonomic database provides for the given taxonomic entity. The list is represented as a string, which is divided with '|' character.

- reference_entity_taxonomic_rank: taxonomic rank of given taxonomic entity.

- reference_entity_phylogenetic_lineage: full taxonomic lineage of given taxonomic entity. Starting from the entity itself on the far left, the parent entity is successively given to the right, ultimately leading to the root of the phylogenetic tree. The list is represented as a string, which is divided with '|' character.

- reference_entity_lineage_taxonomic_rank: taxonomic ranks for each entity within a phylogenetic lineage. 'canonical' taxonomic ranks are given a number greater than 0, while all other ranks are given a number 0. The list is represented as a string, which is divided with '|' character.

- 1: domain, 2: kingdom, 3: phylum, 4: class, 5: order, 6: family, 7: genus, 8: species

● [ref]_genus_dict.txt: [internal_order] - [first_word_block] - [associated_reference_identifier_list]

- internal_order: numerical order of word-blocks (not necessary).

- first_word_block: all word-blocks found within scientific names (canonical names & synonyms) found within a database.

- associated_reference_identifier_list: list of all accession numbers (see reference identifier) which have a preceding word appearing as the first word in one of the associated names. The list is represented as a string, which is divided with '|' character.

If reference metadata is processed into the specified formats and added to the 'pp_ref' directory, PhyloSophos will function normally. 'gbif_preprocessing.py' is an example script which downloads GBIF backbone taxonomy [1] to 'external_files' directory, processes 'Taxon.tsv' file to 'gbif_node_dict.txt' and 'gbif_genus_dict.txt', incorporating GBIF into local PhyloSophos system.

On the other hand, it is possible to remove a particular reference from the 'pp_ref' directory, reducing the number of references PhyloSophos refers to. While doing so reduces the reference import time, it does not significantly affect processing time itself, as it increases the number of input strings to be corrected (status codes >= 20). This also increases the number of 'incorrectly corrected' inputs, such as correcting valid scientific names and mapping them to incorrect DB entries. So we highly discourage to remove taxonomic references from the 'pp_ref' directory.

**Text S5. PhyloSophos modding guide: incorporation of manual curation data**

There is 'manual_curation_list.tsv' file within the /pp_learning directory. If the manual curation option is activated, PhyloSophos imports this file and uses it to convert obsolete synonyms and vernacular name inputs.

Manual curation file has two columns: 'Common_name' column and 'Curated_name' column. The 'Common_name' column contains names that need to be corrected, while the 'Curated_name' column contains the corresponding scientific names that have been manually identified.

You could provide as many [name] - [changed name] pairs as possible, as long as there are no duplicate entries in the 'common_name' column. If there is a duplicate entry, the previous pair of information will be ignored. If the curated name is found to be non-biological (e.g. minerals used in traditional medicine), please enclose the changed name with '<<' and '>>' brackets. PhyloSophos will recognize these flags and return a non-organism flag (mapping status code 90) instead of applying the name input into futile edit-distance mapping steps, thus reducing the processing time.

**Text S6. Case study details & supplementary data files description**

We conducted three case studies to demonstrate the mapping performance of PhyloSophos using data from independent & not primarily taxonomic databases. By doing so, we intended to present a real-life example of how our application can associate detached biological datasets with core references, thus providing a connection to other biological and biochemical information.

All data analysis is based on the taxonomic reference metadata collected on March 8^th^, 2023. We used version 10 (September 16^th^, 2022) metadata of LOTUS initiative [2], and version January 2022 metadata of COCONUT (D-COCONUT) database [3]. We extracted the scientific names included in species-compound pairs, as well as the number of associated compounds per each name, in order to analyze them further. The Compound Combination-Oriented Natural Product Database (K-COCONUT) [4], which was previously available on the internet, is no more accessible now. We listed 8,642 ‘herb’ entries from our previous acquisition of K-COCONUT raw data for further analysis. All three lists of scientific names can be found in supplementary data files.

PhyloSophos analyses were done with the following parameters, which are currently the default setting.

- Reference type: NCBI taxonomy

- Input type: default directory (processing three inputs consequently)

- Levenshtein cutoff: 3

- Manual curation data usage: FALSE

Analysis results could also be found in supplementary data files.

● SD1_phylosophos_d_coconut_analysis.xlsx:

- COCONUT_metadata_processed: processed D-COCONUT raw metadata and chemical association counts

- PhyloSophos_raw_result: raw analysis table produced by PhyloSophos

- Pair_mapping: identification of species-compound pairs which could be associated with valid taxonomic entities (mapping_status_code < 11, singular match).

● SD2_phylosophos_k_coconut_analysis.xlsx:

- K_COCONUT_Raw_species: list of species previously listed on K-COCONUT database

- PhyloSophos_raw_result: raw analysis table produced by PhyloSophos

- Latin_inflections_and_non_bio: individual analysis list of inputs with mapping_status_code 30/31/1000, which consist of inputs with latin inflections or non-biological entities.

● SD3_phylosophos_lotus_analysis.xlsx:

- LOTUS_metadata_processed: processed LOTUS raw metadata and provided mapping information

- PhyloSophos_raw_result: raw analysis table produced by PhyloSophos

- ID_correspondence: comparative analysis between provided mapping information and PhyloSophos-generated mapping information

- NCBI_mapping_disagreement: manual study results about 36 inputs which provided NCBI taxonomic ID is different from PhyloSophos mapping result

- Pair_mapping: identification of species-compound pairs which could be associated with valid taxonomic entities (mapping_status_code < 11, singular match).

**Supplementary figures**

**Supplementary figure 1. Main function of PhyloSophos core mapper script**

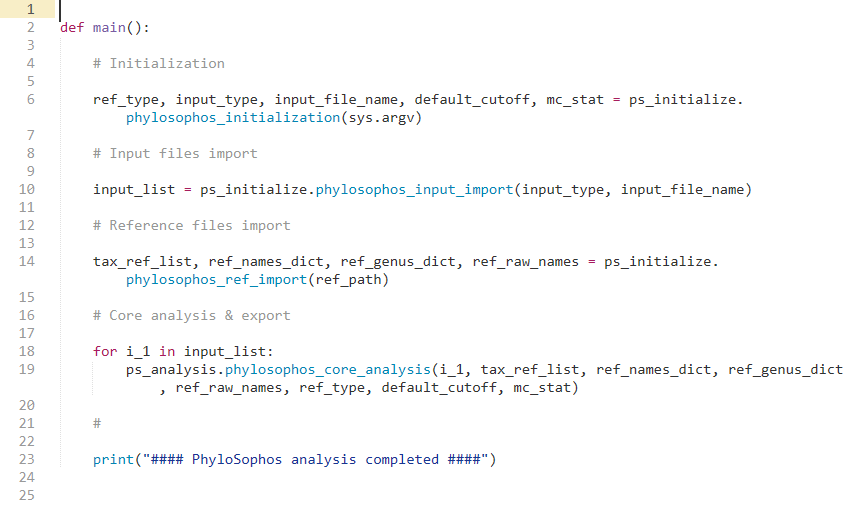

**Supplementary figure 2. Pseudocode of PhyloSophos mapper initialization function**

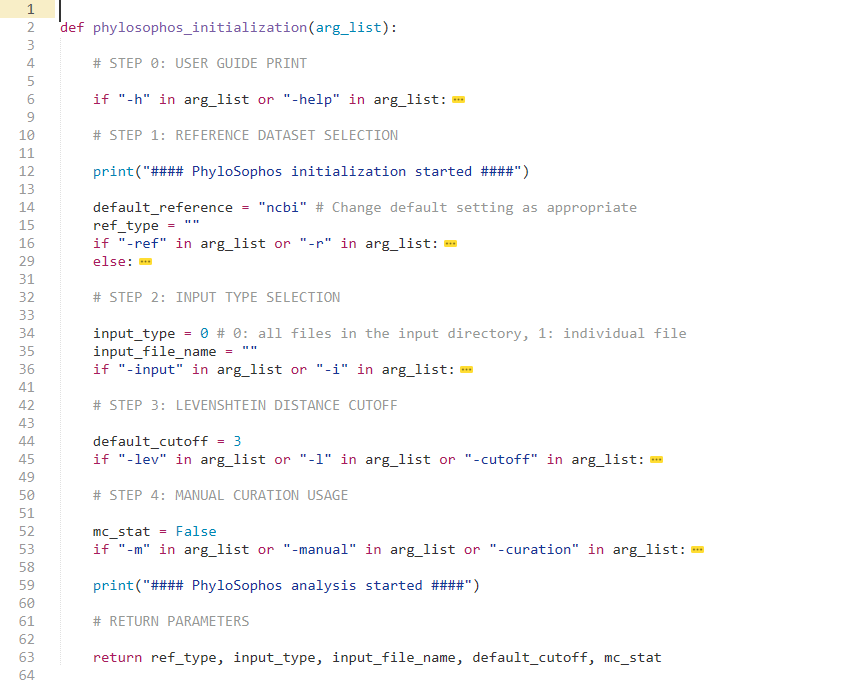

**Supplementary figure 3. Pseudocode of scientific name input import function**

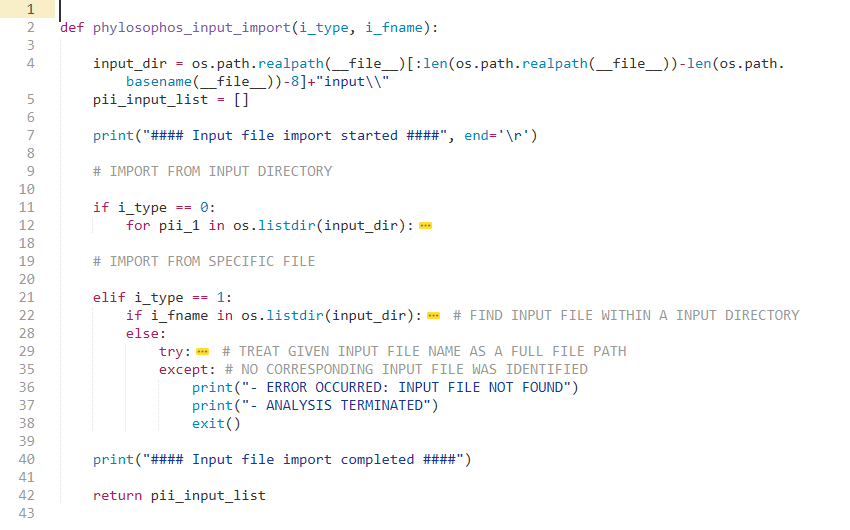

**Supplementary figure 4. Pseudocode of input string-wise PhyloSophos mapping function**

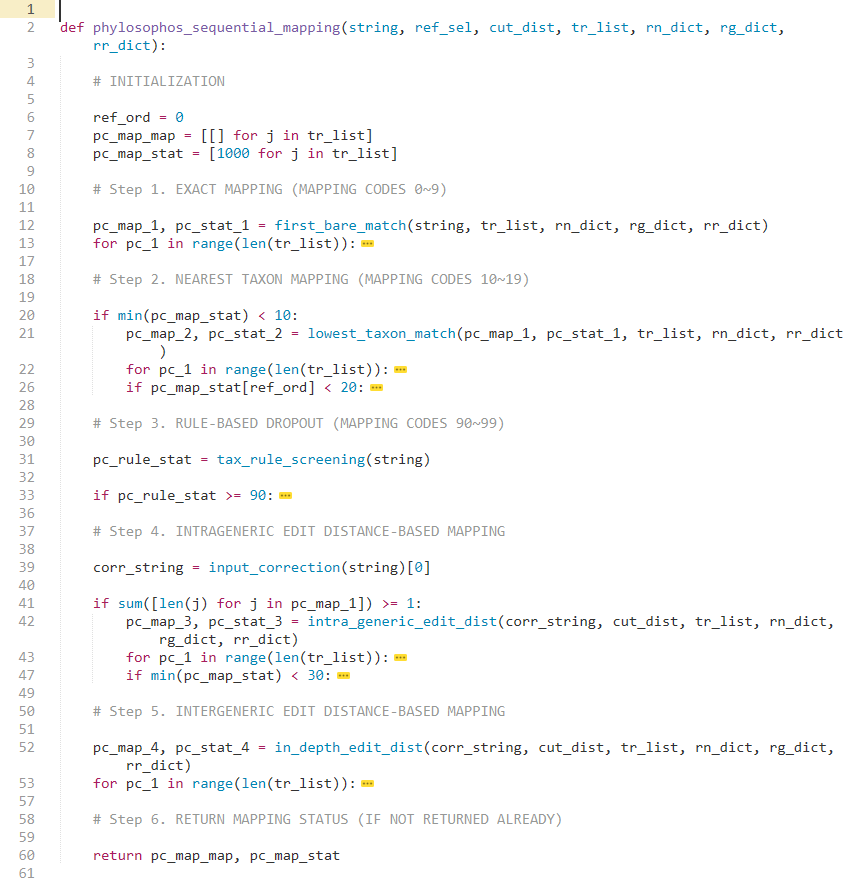

**Supplementary figure 5. Pseudocode of PhyloSophos intrageneric correction subfunction**

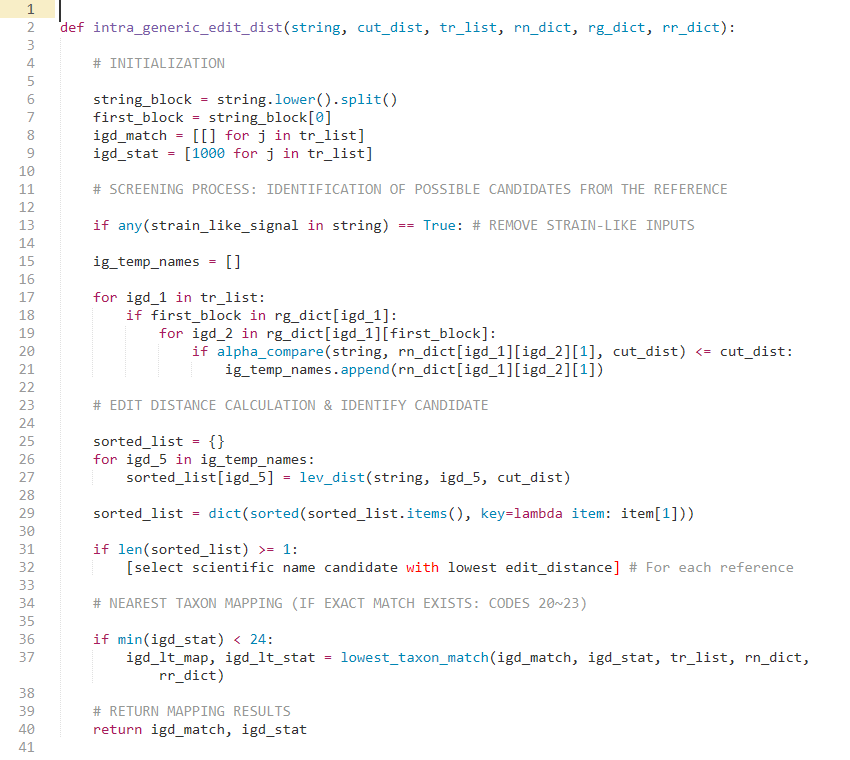

**Supplementary tables**

**Supplementary table 1. PhyloSophos mapping status for scientific names found in natural products databases used in the case study (mapped to NCBI taxonomy)**

| Mapping_category | Mapping_status_code | LOTUS | D-COCONUT | K-COCONUT | Manual_curation_recommendation |
| --- | --- | --- | --- | --- | --- |
| Valid scientific name-Exact match | 0 | 29,109 | 13,353 | 5,980 | NO |
|  | 1 | 2,086 | 2,202 | 333 | NO |
|  | 2 | 11 | 9 | 18 | YES |
|  | 3 | 15 | 1783 | 107 | NO |
|  | 4 | 1 | 158 | 1 | NO |
|  | 5 | 0 | 53 | 0 | YES |
|  | 6 | 704 | 2,088 | 437 | NO |
|  | 8 | 48 | 177 | 46 | YES |
| Valid scientific name-Nearest match | 10 | 256 | 319 | 52 | NO |
|  | 11 | 5,564 | 2,436 | 390 | MAYBE |
|  | 12 | 226 | 50 | 4 | MAYBE |
|  | 13 | 17 | 5 | 1 | MAYBE |
|  | 14 | 0 | 0 | 0 | MAYBE |
|  | 15 | 7 | 2 | 0 | MAYBE |
|  | 16 | 0 | 0 | 0 | MAYBE |
|  | 17 | 0 | 0 | 0 | MAYBE |
| Specific epithet corrected | 20 | 29 | 1,109 | 225 | NO |
|  | 21 | 0 | 0 | 0 | YES |
|  | 22 | 0 | 30 | 1 | NO |
|  | 23 | 0 | 1 | 0 | YES |
|  | 24 | 27 | 266 | 38 | MAYBE |
| Generic/specific epithet corrected | 30 | 0 | 97 | 334 | NO |
|  | 31 | 0 | 0 | 0 | NO |
|  | 32 | 13 | 700 | 140 | NO |
|  | 33 | 0 | 0 | 0 | YES |
|  | 34 | 9 | 302 | 73 | NO |
|  | 35 | 1 | 20 | 7 | YES |
|  | 36 | 0 | 0 | 0 | MAYBE |
| Correction denied | 40 | 0 | 1,712 | 3 | YES |
|  | 41 | 0 | 41 | 1 | YES |
| Mapping denied | 90 | 0 | 0 | 0 | YES |
|  | 91 | 0 | 0 | 0 | YES |
|  | 92 | 0 | 1 | 0 | YES |
|  | 93 | 0 | 0 | 0 | YES |
|  | 94 | 0 | 0 | 0 | YES |
|  | 95 | 0 | 0 | 0 | YES |
|  | 96 | 0 | 12 | 3 | YES |
|  | 97 | 0 | 0 | 12 | YES |
| Partial mapping | 100 | 40 | 1,234 | 173 | YES |
| Unmapped | 1000 | 10 | 368 | 263 | YES |
| (total) | (-) | 38,173 | 28,528 | 8,642 | (-) |

**Supplementary table 2. PhyloSophos broad mapping status (count)**

| Mapping_status_count | LOTUS | D-COCONUT | K-COCONUT |
| --- | --- | --- | --- |
| Valid name / Exact match | 31,915 | 19,823 | 6,922 |
| Valid name / Nearest match | 6,129 | 2,812 | 447 |
| Corrected match | 85 | 2,525 | 818 |
| Partial match | 34 | 2,987 | 177 |
| Dropout | 0 | 13 | 15 |
| Unmatched | 10 | 368 | 263 |
| Total | 38,713 | 28,528 | 8,642 |

**Supplementary table 3. PhyloSophos broad mapping status (ratio)**

| Mapping_status_ratio | LOTUS | D-COCONUT | K-COCONUT |
| --- | --- | --- | --- |
| Valid name / Exact match | 0.836062 | 0.694861 | 0.800972 |
| Valid name / Nearest match | 0.160559 | 0.09857 | 0.051724 |
| Corrected match | 0.002227 | 0.08851 | 0.094654 |
| Partial match | 0.000891 | 0.104704 | 0.020481 |
| Dropout | 0 | 0.000456 | 0.001736 |
| Unmatched | 0.000262 | 0.0129 | 0.030433 |
